# Multi-color live-cell optical nanoscopy using phasor analysis

**DOI:** 10.1101/2023.08.04.551988

**Authors:** Zhimin Zhang, Yuran Huang, Wenli Tao, Yunfei Wei, Liang Xu, Wenwen Gong, Yuhui Zhang, Jiaqiang Zhou, Liangcai Cao, Yong Liu, Yubing Han, Cuifang Kuang, Xu Liu

## Abstract

Stimulated emission depletion microscopy (STED) is a powerful tool for studying nanoscale cell structure and activity, but the difficulties it encounters in multicolor imaging limit its application in biological research. To overcome the disadvantages of limited number of channels and high cost of multicolor STED imaging based on spectral identity, we introduced lifetime into live-cell multicolor STED imaging by separating selected dyes of the same spectrum by phasor analysis. Experimental results show that our method enables live-cell STED imaging with at least 4 colors, enabling observation of cellular activity beyond the diffraction limit.

## Introduction

Fluorescence microscopy is an essential and powerful tool for studying the intricate and dynamic structures inside living cells 1. Over the past two decades, super-resolution fluorescence microscopy (SR) has continued to change the perception of the power of fluorescence microscopy, pushing the resolution of remote optical microscopy to the order of tens of nanometers, allowing the study of nanoscale cell structures under the diffraction limit 2–11. Stimulated emission depletion microscopy (STED) is one of the leading techniques beyond the diffraction limit 3,12. STED has several advantages over other SR methods. Firstly, its immediate super-resolution microscopic properties, i.e. the STED technique does not require any post-processing to achieve super-resolution microscopic imaging, ensuring minimal artifacts 13,14; secondly, in combination with other techniques, such as adaptive imaging methods, it can be applied to deep in vivo imaging 15.

In the last decade, multi-color live cell STED has gained increasing attention and application, most conventionally achieved by using multiple excitation beams and corresponding depletion light 16,17. However, the increasing number of colors not only makes the system more complex and dramatically increasing of the construction cost but also increases the likelihood of photobleaching and more severe phototoxicity, which is not conducive to live cell imaging. Currently, two-color STED is the most commonly achieved with this method 18–21. Besides, fluorescence spectral differentiation is another useful way to realize multicolor STED, e.g. for multicolor STED imaging for emission spectrum light intensity distribution, or multi-channel multicolor imaging using hyperspectral detection 22–24. In addition, other researchers have used methods similar to STORM or PALM to exploit the light-switching properties of fluorescent groups to achieve multi-color STED imaging 25.

In addition to the above-mentioned methods to achieve multi-color structures imaging, one of the important ways to achieve structures segmentation is to exploit the natural difference of the fluorescence lifetime 26–28. In fluorescence lifetime imaging (FLIM), phasor plot analysis is an excellent tool for fluorescence characterization 29. Phasor plot analysis is an amazing analysis method to transform the time-domain features of the fluorescence lifetime into phase-domain features and analyzes its phase features 30. The method is simpler and has clearer fluorescence characteristics as it eliminates the tedious step of fitting curve of the decay data and then obtaining lifetime results for analysis. Structures labeled by different fluorescence probes can be easily characterized using phasor plot analysis. Several researchers have used phase diagram analysis methods to achieve multicolor fluorescence imaging and even multicolor fixed-cell STED imaging 18,31–34. The combination of STED and phasor plot analysis offers fascinating possibilities for multicolor STED imaging of live cells.

To achieve more structural multicolor STED imaging of live cells, we propose multicolor STED live cell imaging based on clustered proportional partitioning of fluorescence lifetime phasor analysis. This method combines the phasor distribution properties of fluorescence with STED super-resolution imaging to achieve more than 3-color STED live cell imaging, culminating in 5-color STED live cell imaging.

## Results

### Multicolor STED based on lifetime identity

Multicolor imaging is essential for studies of organelle interactions, protein co-localization, etc. However, multicolor STED is often costly and difficult to implement because they require the depletion beam wavelength to be at the long trailing end of the fluorescence emission spectrum, and the range of available excitation wavelengths is limited when using a single depletion beam, splitting the limited band into multiple channels will inevitably shorten the detectable fluorescence band and significantly increase the cost of devices such as dichroic mirrors in the system. In addition, finding the right fluorochromes for the right excitation wavelengths and reducing crosstalk between the densely arranged channels is a challenging task. In live cell STED imaging, the limited choice of fluorochromes makes it more difficult to achieve multicolor imaging.

However, wavelength is not the only way to differentiate between fluorescent dyes. The introduction of fluorescence lifetime measurements will add a new dimension to imaging by allowing multiple fluorescent dyes in the same excitation band to be used to label the sample simultaneously, thus avoiding several problems caused by the use of multiple excitation wavelengths. Fluorescence lifetime measurements are achieved by acquiring temporal information about fluorescent photons, which means that no changes to the optical path are required. Fig.1(a-b) demonstrates the segmentation of multiple cellular structures indistinguishable in intensity images by the lifetime identity.

**Fig.1.**
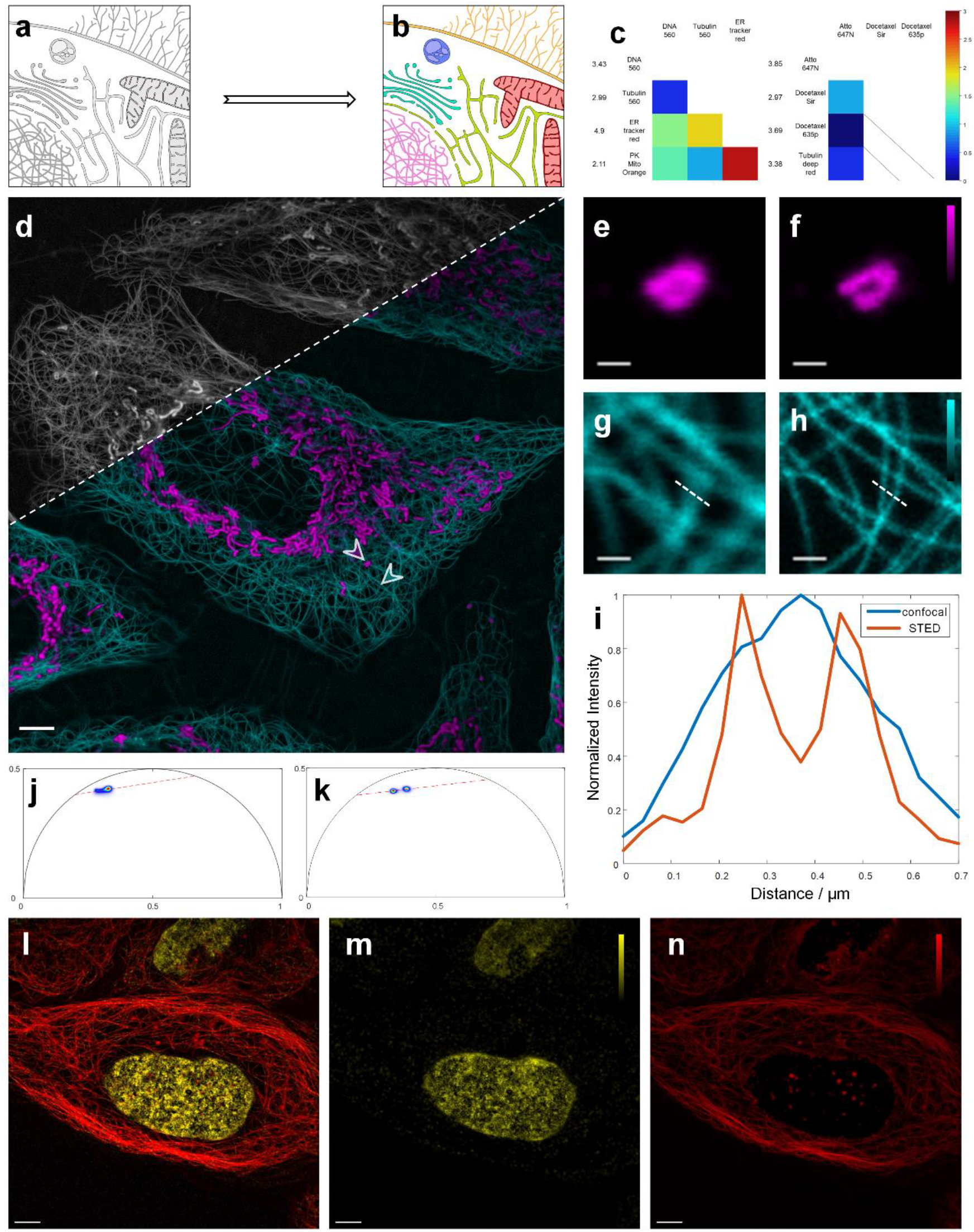
Multiplexing STED based on phasor plot segmentation. **a-b**, Schematic diagram of fluorescence lifetime multiplexing, where six structures (DNA, mitochondria, endoplasmic reticulum, Golgi apparatus, migratory bodies and lysosomes) that could not be distinguished in the intensity map became distinguishable after phasor analysis due to having different fluorescence lifetimes. **c**, Fluorescent dye selection strategy, three commonly used live cell STED dyes were selected at 560 nm and 640 nm excitation wavelength each and the difference in fluorescence lifetimes between them was measured. A larger lifetime difference will lead to better discrimination in phasor analysis, while two dyes with a fluorescence lifetime difference below 0.4 ns are difficult to be separated. **d**, An example of fluorescence lifetime multiplexing, the greyscale image represents the intensity map obtained by STED imaging of live cells, while the colored image demonstrates that mitochondria and microtubules can be discriminated after phasor analysis. These two structures were labeled using DNA560 and ER tracker red, respectively, and were simultaneously excited by a beam at a wavelength of 640 nm. **e-h**, Comparison of STED imaging versus confocal imaging of segmented mitochondria and microtubules, corresponding to the regions indicated by arrows in (d). **i**, Line profile along the dotted lines in (g) and (h), which shows that STED can separate two microtubules in close proximity, revealing details unobservable by confocal images. **j**, The phasor plot corresponding to (d), where the phasor points are projected onto the fitted line and further classified, ultimately leading to the segmentation of the intensity image. **k-n**, another example of fluorescence lifetime multiplexing in wavelength of 560 nm, using TU560 and DNA560 labeling.

Conventional fitting-based fluorescence imaging methods, when applied in multicolor imaging, require the simultaneous fitting of multiple covariates on a pixel-by-pixel basis, which is time-consuming and often gives inaccurate results. The recently developed phasor plot-based method eliminates the need for complex fitting and instead uses a phasor plot to represent the fluorescence lifetime, and the phasor points from pixels with different fluorescence lifetimes are separated from each other in the phasor plot, making it ideal for post-processing calculations for multicolor imaging. By combining the two dimensions of fluorescence lifetime and wavelength, the activity of four biological structures can be observed simultaneously at a resolution beyond the diffraction limit, which is difficult to achieve in other super-resolution methods.

### Imaging methods

As shown in Fig.S1, we built a system consisting of two beams of commonly used pulsed excitation light at wavelengths of 560 nm and 640 nm, and a beam of high power depletion beam at wavelength 775 nm. The fluorescence signal from the sample was received by the avalanche photodiode and its time difference relative to the excitation pulse was acquired by Time-Correlated Single Photon Counting. The total time difference received by each pixel forms a fluorescence decay curve. This curve is determined by the spontaneous radiative excursion rate of the fluorescent molecules and is influenced by the cellular microenvironment, instrument response function, etc. The specific methods for obtaining fluorescence decay and removing the effects of IRF are discussed in Supplementary Note 2.

Following the idea of phasor analysis, the fluorescence decay from each pixel is subsequently transformed into Fourier space to form a series of phasor points. In monochromatic imaging, the phasor points are usually clustered; whereas in our multicolor imaging, the phasor map will consist of several phasor clusters. If the selected fluorescent dyes are too close to each other’s lifetimes, there will be a large overlap between the phasor clusters. Therefore, it is essential to select fluorescent dyes based on their lifetimes. In Fig. 1(c), we selected several commonly used dyes and measured the differences in their lifetime.

If fluorescence saturation is ignored, the fluorescence lifetime is not affected by the excitation light power; however, in sted, the depletion beam forces some fluorescent molecules to return to the ground state by means of excited radiation, which results in the shortening of the fluorescence lifetime, implying that the difference in the lifetimes of the different dyes will also be reduced. Especially troubling is that it also becomes more difficult to separate different structures when we try to improve the resolution by increasing the power of the depletion beam. To address this shortcoming, we added an acousto-optic modulator to the depletion beam path, which enables the depletion beam to switch on and off periodically when the lifetime difference between the dyes is small, allowing the system to switch line by line between the confocal imaging mode and the STED imaging mode, and to use the fluorescence attenuation obtained in the confocal mode as the basis for separating different structures.

### Probes separated by phasor plot

We first show the separation of 2 different structures by phasor analysis, but more structures with sufficiently different fluorescence lifetimes can also be multicolor imaged by the same method. In Figure 1(d) we used Tubulin deep red and Atto 647N to label both microtubules and mitochondria of u2os cells, and because these two dyes have overlapping excitation and emission spectra, they could not be discriminated in STED imaging which only contains intensity identity. However, when the lifetime information was analyzed, we were able to assign different colors to pixels corresponding to different clusters in the phasor plot, when different biological structures were separated. In particular, according to the phasor theorem, pixels with multiple contributions correspond to phasor points located in the middle of several phasor clusters. We project these phasor points onto the line obtained from the fitting, and proportionally distribute the number of photons of the corresponding pixel into the two structures according to the reciprocal of the ratio of their distances to the two phasor clusters. Supplementary Note 3 describes the algorithmic implementation of the above processing. We achieved similar work at 560 wavelengths in Fig.1(k-l), where the significant lifetime difference between DNA560 and TU560 caused 2 distinct clusters to appear in the phasor plot, corresponding to DNA (Fig.1(m)) and microtubules (Fig.1(n)).

Although the lifetime-based multiplexing method is still available in confocal microscopy, its limited resolution may still inconvenience the observation of subtle cellular activities. Here we demonstrate that our multicolor lifetime STED method is capable of simultaneous super-diffraction limited imaging of multiple structures within living cells. Fig.1(e-h) compares the STED versus confocal imaging effects of isolated mitochondria and microtubules, corresponding to the regions indicated by the arrows in Fig.1(d), respectively. The STED images show more details of microtubules and mitochondria, separating the structures that are confounded in the confocal microscope. A more visual comparison is reflected in Figure 1(i), where STED super-resolution produces sharp peaks that are not visible in confocal microscopy.

### Photobleaching compared with normally two-color STED

This section demonstrates that multicolor imaging achieved based on fluorescence lifetimes can reduce photobleaching. In multiplexed STED based on spectral identity, it is common to use different wavelengths of excitation beams to illuminate the sample in turn row by row to avoid severe crosstalk since the fluorescence emission spectra overlap each other. In this case, the depletion beams will scan the sample multiple times, causing severe photobleaching. In Fig. 2 we illustrate that the lifetime-based multiplexed STED method can significantly reduce the damage to the sample from photobleaching due to the fact that the depletion beam only scans the sample a single time. We labeled two sets of u2os cells from the same batch using the same tubulin deep red dye, with the difference that the mitochondria of the first set were labeled using Atto 647N, and the mitochondria and microtubules were excited simultaneously by excitation beams at a wavelength of 640 nm, while those of the second set were labeled using pk mito orange, and samples were scanned with excitation beams at wavelengths of 560 nm and 640 nm, respectively. At 11.4 min of imaging, the brightness of the microtubules in the first set decreases slightly. The statistics shown in Fig.3(c) further indicate that the normalized intensity obtained by the lifetime-based method (red) is consistently higher than that obtained by the spectral-based method (blue).

**Fig.2.**
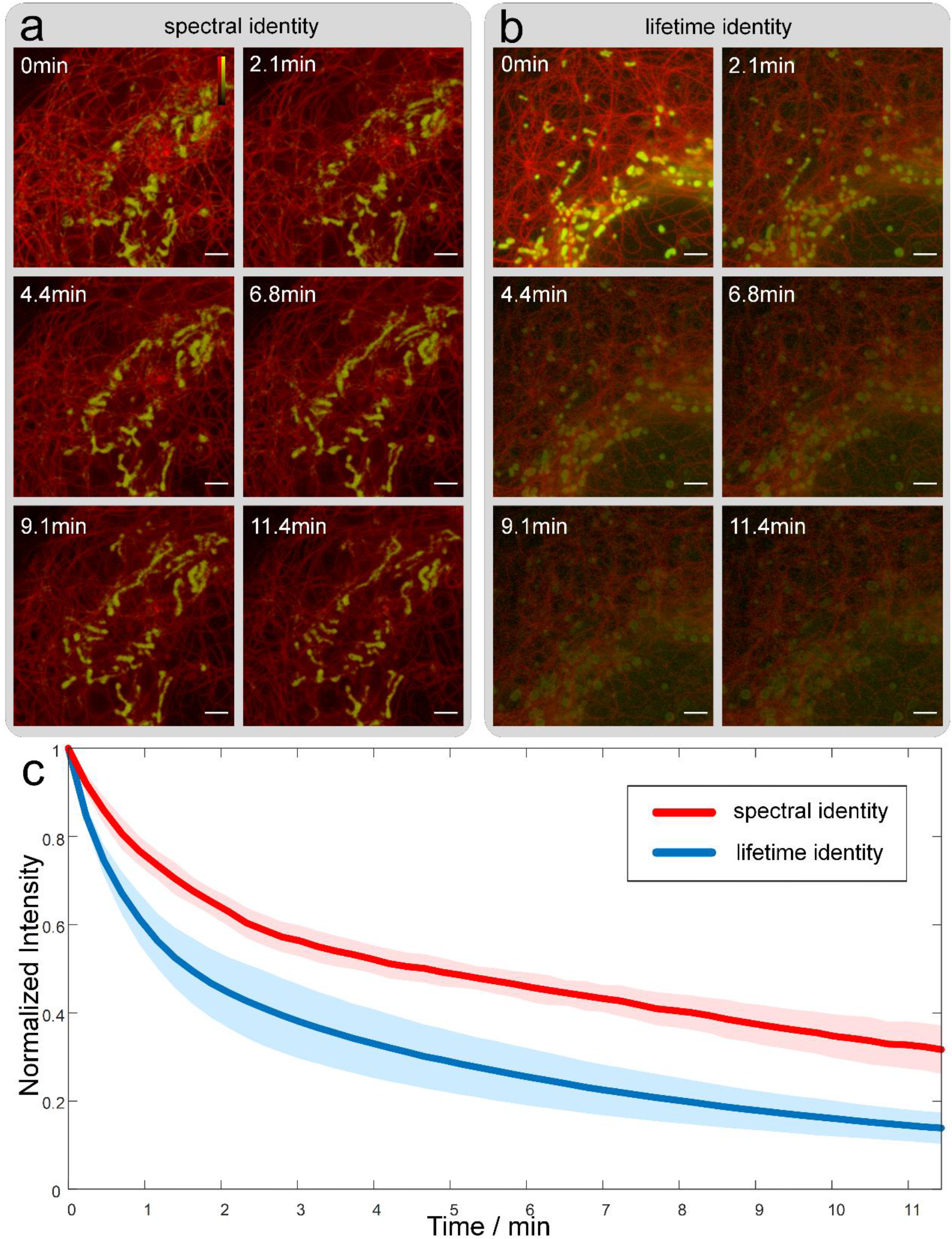
Photostability of lifetime multiplexing STED. **a**, Photostability of mitochondria and microtubules under 11-minute sequential imaging, under the lifetime identity based multiplexing, labeled with Tubulin deep red and Atto 647N and excited simultaneously with a beam at wavelength 640nm. **b**, Photostability under the spectral identity based method, using the same batch of cells that were also labeled with Tubulin deep red, but replacing the mitochondrial dye with pk mito orange, and excited alternately with two beams at wavelengths of 640 nm and 561 nm. **c**, Comparison of photostability of multiplexing STED based on lifetime identity and spectral identity. Lines and shaded areas indicate mean and standard deviation, respectively. The method based on lifetime characteristics has a significantly slower bleaching speed due to the elimination of multiple scans for excitation and depletion beam.

**Fig.3.**
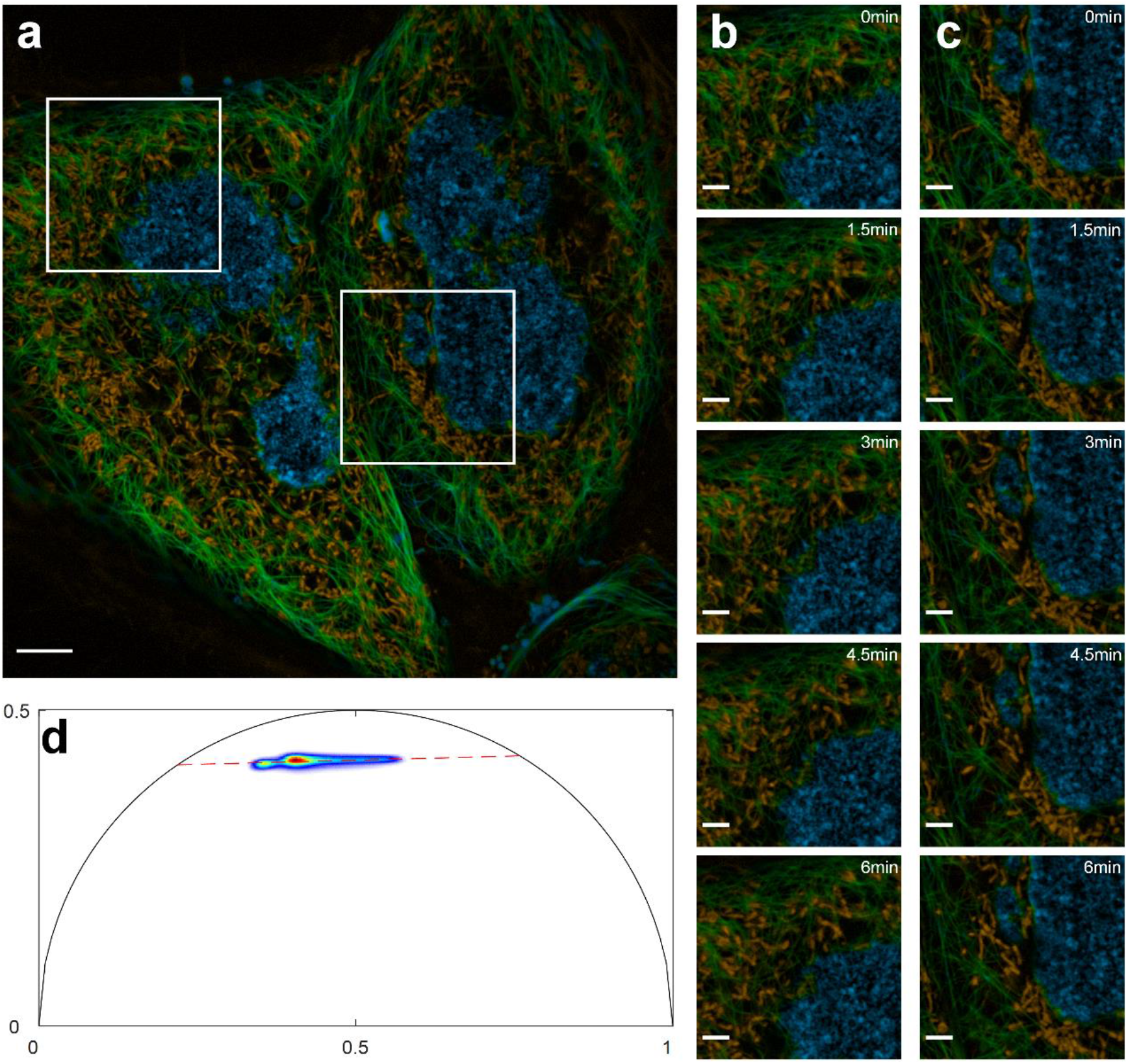
Segmentation of three structures in living cells using phasor analysis. **a**, Three-color imaging results of multiplexed STED method based on lifetime identity, labeled by DNA560, TU560, and pk mito orange **b-c**, Cellular activity in the region marked by the white box in (a) **d**, Corresponding phasor plot

### Multi-color SR imaging of live cells with a single excitation beam

More channels of STED imaging would allow us to simultaneously analyze the interactions of multiple cellular structures with each other, or to look for life activities in which multiple organelles are involved. Expansion of multicolor STED can be achieved by introducing more dyes with different lifetimes at a single excitation wavelength.Fig.3 shows that our analysis is still able to separate the three fluorescent dyes (DNA560, TU560, pk mito orange) that are excited by a beam of light at a wavelength of 560 nm. In pixels with multiple contributions, the distribution of photon numbers still obeys the original law that is inversely proportional to the distance from the phase point to the cluster. After segmentation, the distribution and movement of DNA, microtubules, and mitochondria were efficiently observed.

### Lifetime and spectral based multiplexing STED

Fig.4 demonstrates a feasible 4-color imaging scheme that separates DNA (labeled by DNA560) and endoplasmic reticulum (labeled by er tracker red) under wavelength 560 nm excitation, while separating microtubules (labeled by tubulin deep red) and mitochondria (labeled by Atto 647N) under wavelength 640 nm excitation. We performed long-duration imaging of the four cellular structures and observed mitochondrial division and fusion.

**Fig.4.**
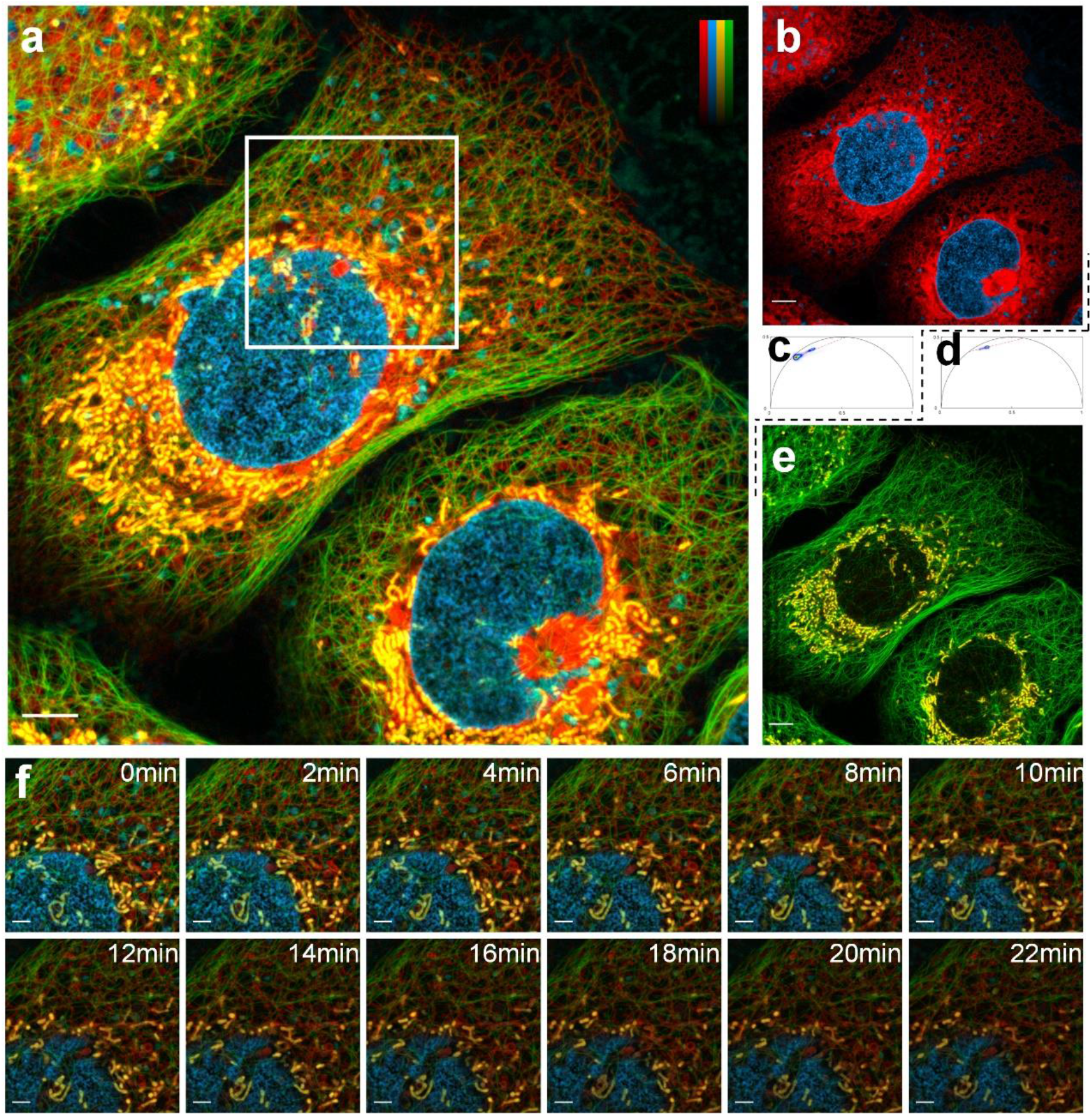
Extending multiplexed STED through spectra and lifetimes.. **a**, Four fluorescent dyes (DNA560, er tracker red, Atto 647N, tubulin deep red) were selected for excitation at wavelengths of 560 nm and 640 nm, respectively, by co-expanding multicolor STED with spectral and lifetime characteristics, and were processed separately in the two excitation channels. **b-c**, imaging results obtained by phasor analysis with excitation at 560 nm and the corresponding phasor plot. **d-e**, imaging results obtained by phasor analysis with excitation at 640 nm and the corresponding phasor plot. **f**, Cellular activity in the boxed region in (a) during 22 minutes

## Discussion

Achieving multicolor STED in living cells is very difficult and costly due to the limitation of the depletion beam on the excitation spectrum. However, phasor plot-based fluorescence lifetime imaging opens up new possibilities for the realization of multicolor STEDs in living cells. By selecting a suitable STED and analyzing it with high precision, we have demonstrated that 3-color imaging is possible using a single wavelength of excitation light and that 4 different structures can be distinguished simultaneously when the excitation light is extended to two different wavelengths. We believe that the advantages of high-resolution, high-sensitivity, and multi-structure lifetime based multicolor STED can facilitate the study of cell metabolism, organelle interactions, protein co-localization, and other biological topics.

Theoretically, there is no limit to the number of structures in lifespan-based multicolor STEDs; however, as the number of structures increases, finding the corresponding fluorescent dyes with sufficient lifetime differences becomes difficult, (staining difficulties). In our experience, two fluorescent dyes with a lifetime difference of more than 0.4 ns can be well separated, but this is still subject to the actual imaging conditions such as the microenvironment in which the dye molecules are embedded. Overall. Achieving more structural STED imaging of living cells will depend on the discovery of more dye combinations.

## Acknowledgment

We thank Z.Y.Liu for his help with imaging processing.

This work was supported by the following grants: National Natural Science Foundation of China (61827825, 62125504); STI 2030—

Major Projects+2021ZD0200401; China Postdoctoral Science Foundation(BX2021272); Zhejiang Provincial Ten Thousand Plan for Young Top Talents (2020R52001).

## Author contributions

Conceived and oversaw the project: Y.B.H, C.F.K, Y.H.Z. Initiated the designs for microscope and imaging method: C.F.K. Initiated the designs for biological experiments: Y.B.H, Y.H.Z. Designed and developed the microscope: Z.M.Z, Y.R.H, L.X. Designed the imaging software: Z. M.Z. Designed the processing algorithm: Y.R.H, Z.M.Z. Prepared the experimental samples: W.L.T, W.W.G. Synthesized the fluorescent probes: Y.F.W. Performed the experiments: Y.R.H, W.L.T. All authors inspected data and contributed to the drafting of the manuscript. Supervi sed resarch: Y.B.H, C.F.K, Y.H.Z, X.L. Directed research: Y.B.H.

